# A novel, tissue-associated and vertically transmitted bacterial symbiont in the coral *Pocillopora acuta*

**DOI:** 10.1101/2023.09.14.557667

**Authors:** Justin Maire, Sarah Jane Tsang Min Ching, Katarina Damjanovic, Hannah E. Epstein, Louise M. Judd, Linda L. Blackall, Madeleine J. H. van Oppen

**Affiliations:** School of BioSciences, The University of Melbourne, Parkville, 3010, VIC, Australia; Australian Institute of Marine Science, PMB No 3, Townsville, 4810, QLD, Australia; ARC Centre of Excellence for Coral Reef Studies, James Cook University, Townsville, 4811, QLD, Australia; Doherty Applied Microbial Genomics, Department of Microbiology and Immunology, The University of Melbourne at the Peter Doherty Institute for Infection and Immunity, Parkville, 3010, VIC, Australia

**Keywords:** *Endozoicomonas*, endozoicomonadaceae, cnidarian, symbiosis, microbiome, genomics

## Abstract

Coral microhabitats are colonized by a myriad of microorganisms, including diverse bacteria which are essential for host functioning and survival. However, the location, transmission, and functions of individual bacterial species living inside the coral tissues remain poorly studied. Here, we show that a previously undescribed bacterial symbiont of the coral *Pocillopora acuta* forms cell-associated microbial aggregates (CAMAs) within the mesenterial filaments. CAMAs were found in both adults and larval offspring, providing evidence of vertical transmission. *In situ* laser capture microdissection of CAMAs followed by 16S rRNA gene metabarcoding and shotgun metagenomics produced a near complete metagenome-assembled genome. We subsequently cultured the CAMA bacteria from *P. acuta* colonies, and sequenced and assembled their genomes. Phylogenetic analyses showed that the CAMA bacteria belong to an undescribed Endozoicomonadaceae genus and species, which we propose to name *Sororendozoicomonas aggregata* gen. nov sp. nov. Metabolic pathway reconstruction from its genome sequence suggests this species can synthesize most amino acids, several B vitamins, and antioxidants, which may be beneficial to its coral hosts. This study provides detailed insights into a new member of the widespread Endozoicomonadaceae family, thereby improving our understanding of coral holobiont functioning. Vertically transmitted, tissue-associated bacteria, such as *S. aggregata* may be key candidates for the development of microbiome manipulation approaches with long-term positive effects on the coral host.

## Main text

Corals associate with diverse bacteria with essential functions, such as protection against pathogens and nutrient cycling [1]. Among these bacteria, a small portion resides within the coral tissues [2], sometimes forming cell-associated microbial aggregates (CAMAs) [3–6]. However, the identification and characterization of CAMA-forming bacteria in corals remains in its infancy. The most common CAMA bacteria belong to the *Endozoicomonas* genus [3, 4, 7], widespread coral symbionts generally considered to be mutualistic through sulfur cycling and B vitamin synthesis [3, 7–10]. *Simkania* sp. (Chlamydiota) was recently shown to also form CAMAs in *Pocillopora acuta* from Feather Reef (Great Barrier Reef [GBR], Australia; Figure S1) [3], although its function remains unclear.

Here, we characterized CAMAs present in *P. acuta* colonies from Orpheus Island, also in the GBR. *P. acuta* is an asexual brooder, producing fully formed, genetically identical larvae, and a sexual broadcast spawner [11]. The microbiome of Orpheus Island colonies is dominated by *Endozoicomonas* in early and adult life stages [6, 12], and CAMAs were detected in larvae, indicating possible vertical transmission [6]. Using samples from these two studies (Figure S2A-B), we verified by fluorescence *in situ* hybridization (FISH) that both adults (X7 colony [12]) and larvae (OI2 and OI3 colonies [6]) possessed CAMAs (Figure 1A-E). In adult polyps, CAMAs localized to the mesenterial filaments (Figure 1A-C). In larvae, CAMAs were found in the gastrodermis (Figure 1D-E).

**Figure 1:**
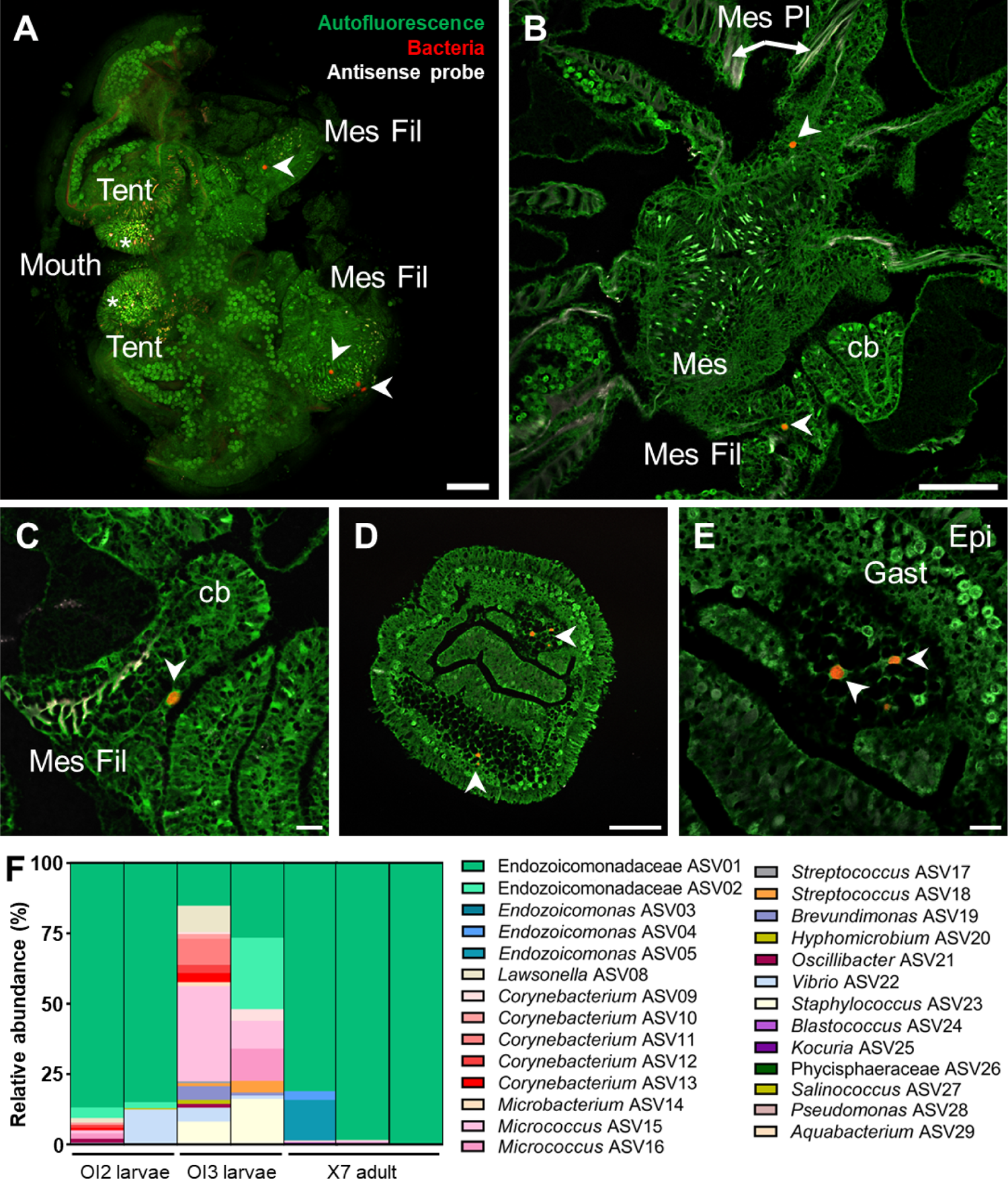
Location and identity of CAMAs in Orpheus Island *Pocillopora acuta*. A: CAMAs in an adult polyp (X7 colony) by whole-mount FISH. B-E: FISH images showing the location of CAMAs in sectioned adult polyps (B, C; X7 colony) and sectioned larvae (D, E; OI2 colony). White arrowheads point at CAMAs. Green: autofluorescence; red: EUB338-mix probe (all bacteria); white: non-EUB probe (negative control). Note the non-specific binding in tentacles (red and white signal overlapping, white asterisks), often localized in nematocysts. Tent: tentacle; Mes Fil: mesenterial filaments; Mes: mesenteries; Mes Pl: mesogleal plates; cb: cnidoglandular band; Gast: gastrodermis; Epi: epidermis. Scale bars: 100 µm for A, B, D; 20 µm for C and E. F: Relative abundance of bacterial ASVs in CAMAs of larvae (OI2 and OI3 colonies) and adult branches (X7 colony) isolated by LCM. Each bar represents one replicate (either three pooled larvae or one adult branch). Detailed composition is available in Table S2.

To taxonomically identify the bacteria present in the CAMAs, we excised CAMAs using laser capture microdissection (LCM), followed by 16S rRNA gene metabarcoding (Figure S3, Table S1). Two ASVs assigned to *Kistimonas* (Endozoicomonadaceae ASVs 01 and 02) made up most of the reads in OI2 larvae and X7 branches (Figure 1F, Table S2). While the samples were collected at the same site, they originated from different host colonies and were collected several months apart, highlighting the temporal stability of this association among individuals and through ontogeny. *Kistimonas* is part of the Endozoicomonadaceae family along with *Endozoicomonas*. Because of taxonomic discrepancies discussed below, we refer to these two ASVs as Endozoicomonadaceae ASVs 01 and 02. The two ASVs were also present in OI3 larvae at a lower relative abundance. In addition, three *Endozoicomonas* ASVs were found in X7 and were identical to *Endozoicomonas* present in *P. acuta* from Feather Reef [3]. FISH using an Endozoicomonadaceae probe provided additional support for this taxonomy in both adult and larval CAMAs (Figure S4).

While *Kistimonas* was not reported in the original studies [6, 12], a re-analysis of both previous studies reassigned three *Endozoicomonas* ASVs to *Kistimonas* (Figure S5, Table S3), two of which were 100% identical to Endozoicomonadaceae ASVs 01 and 02 found in our excised CAMA samples. Virtually no *Endozoicomonas* strains (only one ASV at 0.02-0.04% relative abundance) were detected in whole larvae or recruits in our re-analysis of the earlier datasets (Figure S5), suggesting that *Endozoicomonas* are acquired horizontally in Orpheus Island *P. acuta*, as observed for Feather Reef *P. acuta* [3]. Conversely, Endozoicomonadaceae ASVs 01 and 02 were present at all life stages (*i.e.* larvae, recruits and adults of the same colonies, Figure S5) and formed CAMAs, confirming their vertical transmission. Whether whole CAMAs or individual bacteria are transmitted from parent colony to asexually produced larvae remains unknown. Strong evidence for vertical transmission of bacteria in corals has only been reported twice before [13, 14].

Shotgun sequencing of the X7 CAMA samples yielded one metagenome-assembled genome (MAG) (Pac_X7, Table S4), the 16S rRNA gene sequence of which was identical to Endozoicomonadaceae ASV01. An additional *P. acuta* colony (P3, Figure S2C) was collected from Orpheus Island in June 2023, from which we cultured ten bacterial isolates with 16S rRNA gene sequences identical to Pac_X7, one of which was whole-genome sequenced (Pac_P3-11-1, Table S4). Both assemblies had 99.8% Average Nucleotide Identity (ANI) and 99.6% Average Amino acid Identity (AAI) (Figure S6), confirming that the Pac_P3-11-1 isolate is indeed a CAMA bacterium. Phylogenetic analysis of 120 marker genes placed both genomes separately from all described Endozoicomonadaceae genera (Figure 2A, Table S5), along with Endozoicomonadaceae SCSIO12664 (99.6% 16S rRNA gene sequence identity, 99.2% ANI, 98.8% AAI), isolated from *Pocillopora damicornis* in the South China Sea [15]. This was supported by 16S rRNA gene phylogeny (Figure S7). Together with low AAI (highest score of 60.1% with *E. elysicola*, Figure S6) and low 16S rRNA gene identity (highest score of 93.97% with *E. gorgoniicola*), these data support the placement of Pac_P3-11-1 and Endozoicomonadaceae SCSIO12664 in a new genus and species, which we propose to name *Sororendozoicomonas aggregata*, gen. nov., sp. nov. This likely explains the discrepancies observed with metabarcoding-based taxonomy, as taxonomy would be assigned to a closely related taxon rather than an undescribed genus.

**Figure 2:**
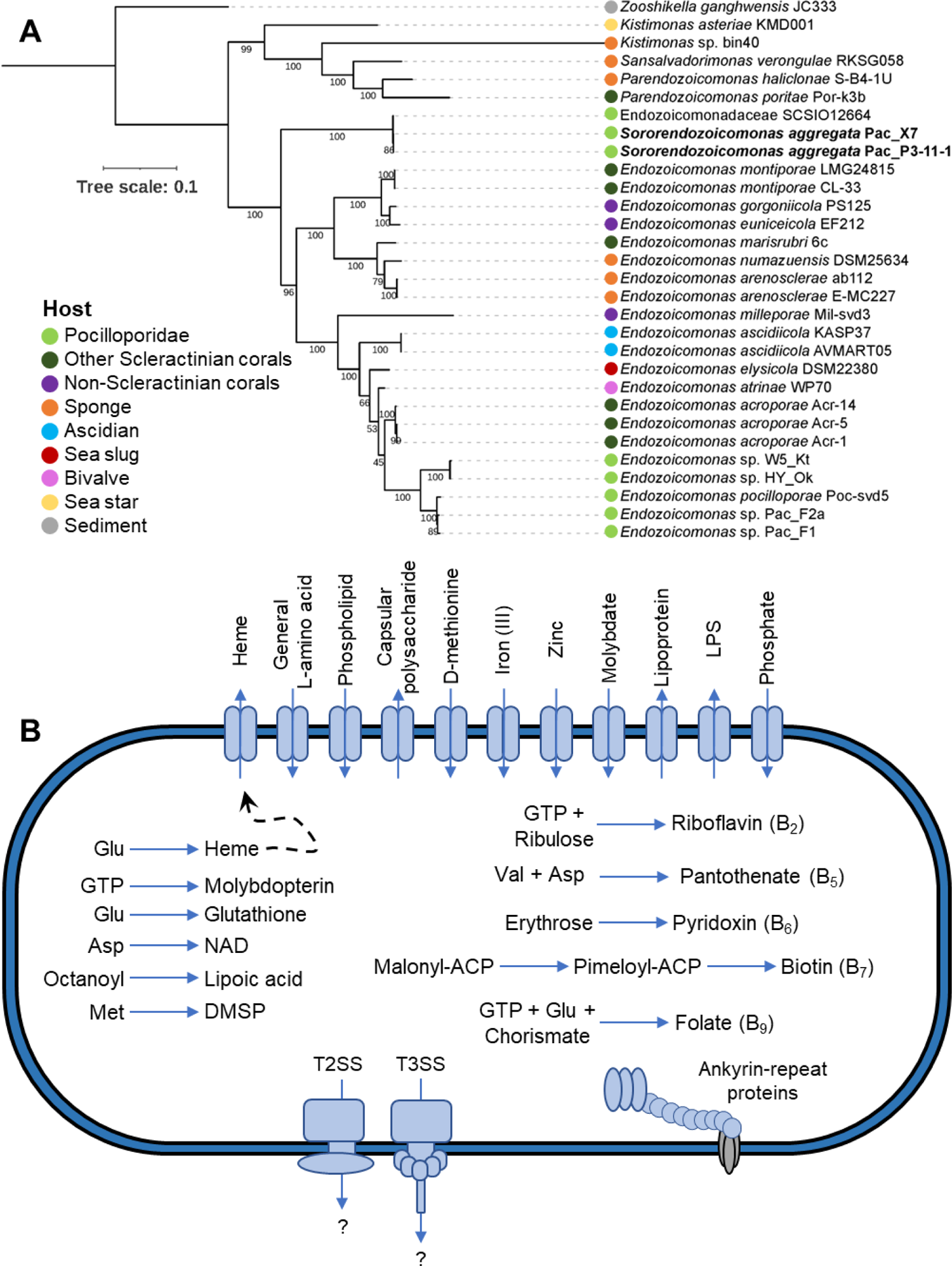
Taxonomy and functional potential of *Sororendozoicomonas aggregata*, CAMA member of Orpheus Island *Pocillopora acuta*. A: Maximum likelihood phylogeny of Endozoicomonadaceae based on 120 marker genes and 28 Endozoicomonadaceae genomes in addition to the two genomes recovered in this study. *Zooshikella* was chosen as the outgroup. Bootstrap support values based on 1000 replications are provided. Additional data on the reference genomes are available in Table S5. B: Overview of the metabolic potential from the genome sequence of Pac_P3-11-1 recovered from *P. acuta* CAMAs. The pathways represented here are at least 80% complete (Fig. S7). Dashed arrows represent hypotheticals. T2SS: type II secretion system; T3SS: type III secretion system; DMSP: dimethylsulfoniopropionate; LPS: lipopolysaccharide; NAD: nicotinamide adenine dinucleotide.

Genome annotation (Figures 2B and S8, Table S6) revealed Pac_P3-11-1 encodes several genes indicative of host-symbiont interactions, including type II and type III secretion systems, 35 eukaryotic-like proteins (Table S7), and three secondary metabolites with putative antimicrobial activity (Table S8). This gene repertoire is similar to that of other coral-associated Endozoicomonadaceae [3, 4, 8–10] and may regulate host colonization, aggregation, and vertical transmission. Additionally, we recovered pathways for the biosynthesis of all amino acids (except phenylalanine, tyrosine, and arginine), as well as riboflavin (vitamin B_2_), pantothenate (B_5_), pyridoxin (B_6_), biotin (B_7_), and folate (B_9_), which could assist with coral metabolism [3, 4, 9, 10, 16]. Finally, Pac_P3-11-1 shows potential for the scavenging of reactive oxygen species, which are believed to drive coral bleaching when present in excess [17], as it possesses pathways for the synthesis of the antioxidants heme, lipoic acid, and glutathione, and the *dsyB* gene, essential for the synthesis of dimethylsulfonioproprionate [18].

In conclusion, we described and cultured the first coral tissue-associated, vertically transmitted member of the Endozoicomonadaceae family. *Sororendozoicomonas* was not found in other GBR *P. acuta* populations that harbor *Endozoicomonas* [3], suggesting geographical location may impact Endozoicomonadaceae associations. Additionally, *Endozoicomonas* CAMAs were previously found in pocilloporid tentacles [3, 4], contrasting with *Sororendozoicomonas*’ location in the mesenterial filaments.

As climate change increasingly triggers coral bleaching and mortality, novel methods are urgently needed to increase coral climate resilience, including the manipulation of coral-associated microorganisms. Endozoicomonadaceae are great candidates for coral microbiome manipulation approaches [19, 20], due to their prevalence, tissue location (*i.e.* more stably associated than bacteria residing in the mucus or skeleton), and putative benefits. The vertical transmission and cultivability of *S. aggregata* make it an ideal candidate, ensuring its stability within the coral holobiont across generations and thereby the long-term sustainability of conservation/restoration approaches based on microbiome manipulation.

### Description of *Sororendozoicomonas* gen. nov

*Sororendozoicomonas* (So.ror.en.do.zo.i.co.mo’nas. L. fem. n. soror, sister; *Endozoicomonas*, taxonomic name of a bacterial genus; *Sororendozoicomonas* referring to the close relationship to the bacterial genus *Endozoicomonas*). The name was registered through SeqCode [21].

The genus *Sororendozoicomonas* represents a distinct lineage within the Endozoicomonadaceae family, sister to *Endozoicomonas*, supported by concatenated marker gene phylogenies (Figures 2A) as well as 16S rRNA gene phylogeny (Figure S7) in the present study. *Sororendozoicomonas* show similar functional and metabolic potential to *Endozoiocomonas* species (Figure 2B). The new genus contains two strains isolated from *Pocillopora* corals (Pac-P3-11-1, from the present study, and SCSIO12664 [15]) and potentially additional strains isolated from Red Sea *Stylophora pistillata* [22] (only 16S rRNA gene sequences are available - KC669256.1, KC669230.1, KC669217.1, KC669131.1).

### Description of Sororendozoicomonas aggregata sp. nov

*Sororendozoicomonas aggregata* (ag.gre.ga’ta. L. fem. part. adj. *aggregata,* joined together, referring to the ability to form aggregates within coral tissues). The name was registered through SeqCode [21]. The new species contains two strains isolated from *Pocillopora* corals (Pac-P3-11-1, from the present study, and SCSIO12664 [15]).

Type strain: Pac_P3-11-1, isolated from tissues of the coral *Pocillopora acuta*, collected from Orpheus Island, Australia (18°36’16.26“S, 146°29’24.27” E). Taxonomic assignment based on concatenated marker gene phylogenies (Figure 2A) as well as 16S rRNA gene phylogeny (Figure S7). Grows on Marine Agar 2216 (Difco) between 23°C and 28°C, forming circular, white, raised, sticky colonies with entire margins.

## Acknowledgements

This research was supported by the Australian Research Council (FL180100036 to MJHvO; DP160101468 to MJHvO and LLB), the Native Australian Animals Trust (to JM), Paul G. Allen Philanthropies (to MJHvO), and AIMS@JCU, the joint venture between James Cook University and the Australian Institute of Marine Science, as part of H.E.E’s PhD research. H.E.E acknowledges receipt of an AIMS@JCU Postgraduate Scholarship. We thank: Corinne E. Allen for the collection of coral colonies from Orpheus Island (permit G22/46479.1); the Biosciences Microscopy Unit (University of Melbourne) and the Biological Optical Microscopy Platform (University of Melbourne) for the use of their confocal microscopes and laser capture dissection, and particularly Dr Gabriela Segal, Dr Shane Cheung and Dr Sam Mills for their valuable assistance; the Melbourne Histology Platform (University of Melbourne), Laura Leone, and Lisa Foster for their assistance with sample sectioning; Dr Kshitij Tandon and Dr Gayle K. Philip for assistance with genome analyses; the Melbourne Research Cloud, the University of Melbourne’s Research Computing Services and the Petascale Campus Initiative for providing the high-performance computing instances needed for this work.

## Data availability statement

Raw data are available under NCBI BioProject IDs PRJNA891898 (larvae CAMAs, MiSeq raw data) and PRJNA974967 (adult CAMAs, MiSeq and NovaSeq raw data, Pac_X7 MAG assembly; Pac_P3-11-1 NextSeq raw reads and assembly). The complete 16S rRNA gene sequence of Pac_P3-11-1 is available under the accession number OR505856.

## Author contribution

Conceptualization: JM, MvO, LB; Investigation: JM, SJTMC, KD, HE, LJ; Methodology: JM; Resources: KD, HE; Formal analysis: JM; Visualization: JM; Funding acquisition: MvO, LB, JM; Writing - Original Draft Preparation: JM, MvO. All authors reviewed and edited the final manuscript.

## Material and methods

### Coral larvae collection (OI2 and OI3 colonies)

For OI2 and OI3, parent colonies were collected in 2017 from Orpheus Island (Little Pioneer Bay; 18°36’3.6“S 146°29’20.4”E), in the central Great Barrier Reef in Australia (Figure S1 and S2A), and maintained and sampled as part of a previous experiment [6]. Briefly, before planulation, colonies were maintained in individual acrylic aquaria that received indirect natural sunlight and 0.4 μm-filtered seawater. A filter was fitted at each outlet of the acrylic tank to collect released planulae. Larvae were washed with 0.22-µm filtered seawater (FSW), transferred from the filter into a microcentrifuge tube using a sterile pipette tip, and as much water as possible was removed from the tube without disturbing the larvae. Larvae were fixed for 10 hrs at 4°C in 4% paraformaldehyde (PFA) prepared in FSW, rinsed twice in FSW, and stored in 50% ethanol-PBS at −20°C. Around 10 larvae per parent colony were fixed.

### Adult coral collection (X7 and P3 colonies)

For the X7 colony (Figure S2B), coral fragments were sampled from the field in 2016 from the same site at Orpheus Island as part of a previous experiment (Figure S1) [12]. From this study, only samples collected in November 2016 were processed here. Four small coral fragments (2-5 cm in length) were snapped off the colonies with forceps, fixed for 24 hrs in 4% PFA in FSW, rinsed twice in FSW, and stored in 50% ethanol-PBS at −20°C. Following fixation, coral branches were decalcified in EDTA 10%. EDTA was renewed every two days, and samples were kept at 4°C on a rotating wheel, until there was no skeleton left (around two weeks). Samples were then rinsed in PBS 1X, and stored at 4°C in PBS 1X.

For the P3 colony (Figure S2C), coral colonies were collected at 1-4 m depth with a chisel and hammer from Orpheus Island (Little Pioneer Bay, 18°36’16.26“ S, 146°29’24.27” E, Collection permit G22/46479.1) in June 2023. Four colonies were brought back to the Australian Institute of Marine Science for overnight holding before being shipped to the University of Melbourne the next day. On arrival, the corals were placed in 130 L recirculating system tanks and kept there overnight for 10 h at 25 °C in 35 p.p.t. reconstituted sea water (Red Sea Salt^TM^, R11065, Red Sea, USA).

### Fluorescence *in situ* Hybridization (sections)

Sample processing, embedding, and sectioning (3-µm thickness) were performed by the Melbourne Histology Platform (University of Melbourne) as previously described [3]. Fluorescence *in situ* hybridization (FISH) was then performed as previously described [6], but final probe concentration during hybridization was 5 ng/µL. See Table S9 for probe sequences, fluorophores, and formamide concentrations. Slides were mounted in CitiFluor^TM^ CFM3 mounting medium (proSciTech, Australia), covered with a coverslip and sealed with clear nail polish. Slides were kept at 4°C until observation.

### Fluorescence *in situ* Hybridization (whole-mount)

Single polyps of the X7 colony were dissected in PBS 1X using Dumont tweezers under a dissecting microscope. Samples were cleared of autofluorescence by incubating polyps in methanol-PBS 1X (50:50 v/v) for 10 min, methanol-PBS 1X (75:25 v/v) for 10 min, methanol-PBS 1X (90:10 v/v) for 10 min, 100% methanol for 10 min, methanol-PBS 1X + Triton X-100 0.2% (90:10 v/v) for 10 min, methanol-PBS 1X + Triton X-100 0.2% (75:25 v/v) for 10 min, methanol-PBS 1X + Triton X-100 0.2% (50:50 v/v) for 10 min, PBS 1X + 0.2% Triton X-100 for 10 min, and stored at 4°C in PBS 1X. FISH was performed on whole polyps as previously described [3], with the probes shown in Table S9. Before observation, single polyps were cut in half to expose the gastric cavity and mesenteries, and deposited onto 8-well coverslip-bottom slides (ibidi, USA) with a drop of milliQ water to avoid drying.

### Confocal Laser Scanning Microscopy

Slides were observed on a Nikon A1R confocal laser scanning microscope (Nikon, Japan) with the NIS325 Element software. Virtual band mode was used to acquire variable emission bandwidth to tailor acquisition for specific fluorophores. The fluorophores Atto550 were excited using the 561 nm laser line, Atto647 using the 640 nm laser line, and the coral autofluorescence using the 488 nm laser line with a detection range of 570-620 nm for Atto550, 660-710 nm for Atto647, and 500-550 nm for coral autofluorescence. For three-dimensional reconstructions of Z-stacks (for whole polyps), sections were acquired using Z steps of 3.2 μM with the 10X objective. Nd2 files were processed using ImageJ. Z-stacks were projected in two-dimensional images using the ‘Max Intensity’ projection type. Linear adjustments of brightness and contrast were performed when necessary and applied to the entire image and to each channel independently. Channels were then given artificial colors (see figure legends) and merged.

### Laser Capture Microdissection of CAMAs and DNA extraction

Laser capture microdissection (LCM) of CAMAs in OI2, OI3, and X7 samples was performed as previously described [3]. For OI2 and OI3 larvae, three replicates each containing three larvae were processed. For each replicate, ten slides each containing eight sections were processed (∼80 3-µm sections per replicate in total). For X7 branches, three replicates each containing one branch were processed. For each replicate, 15 slides each containing four sections were processed (∼60 3-µm sections per replicate in total). Tissue areas without CAMAs were also separately captured as a negative control. DNA extraction was then conducted as previously described [3], using the Arcturus® PicoPure® DNA Extraction Kit (Applied Biosystems, USA). Three caps containing no captured tissue, but that were open in the LCM facility to capture air contamination, were also included as extraction blanks.

### 16S rRNA gene metabarcoding

Hypervariable regions V5-V6 of the 16S rRNA genes were amplified using the primer set 784F (5ʹ <u>GTGACCTATGAACTCAGGAGTC</u>AGGATTAGATACCCTGGTA 3ʹ) and 1061R (5ʹ <u>CTGAGACTTGCACATCGCAGC</u>CRRCACGAGCTGACGAC 3ʹ). Adapters were attached to the primers and are underlined. Bacterial 16S rRNA genes were PCR-amplified on a SimpliAmp Thermal Cycler (Applied Biosystems, ThermoFisher Scientific). Each reaction contained 1 μL of DNA template, 1.5 μL of forward primer (10 μM stock), 1.5 μL of reverse primer (10 μM stock), 7.5 μL of 2x QIAGEN Multiplex PCR Master Mix (Qiagen, Germany) and 3.5 μL of nuclease-free water (Thermofisher), with a total volume of 15 μL per reaction. Three triplicate PCRs were conducted for each sample and three no-template PCRs were conducted as negative controls. PCR conditions for the 16S rRNA genes were as follows: initial denaturation at 95°C for 3 min, then 18 cycles of: denaturation at 95^°^C for 15 s, annealing at 55^°^C for 30 s, and extension at 72^°^C for 30 s; with a final extension at 72^°^C for 7 minutes. Samples were then held at 4^°^C. Following PCR, triplicates were pooled, resulting in 45 μL per sample. Metabarcoding library preparation was conducted as previously described [23] and sequencing was performed at the Walter and Eliza Hall Institute (WEHI) in Melbourne, Australia on one MiSeq V3 system (Illumina) with 2×300bp paired-end reads.

### 16S rRNA gene metabarcoding analysis

QIIME2 v 2021.8 [24] was used for processing 16S rRNA gene sequences. The plugin demux [24] was used to create an interactive plot to visualize the data and assess the quality, for demultiplexing and quality filtering of raw sequences. The plugin cutadapt [25] was used to remove the primers and MiSeq adapters. DADA2 [26] was used for denoising and chimera checking, trimming, dereplication, generation of a feature table, joining of paired-end reads, correcting sequencing errors, and removing low quality reads (Q-score < 30). Summary statistics were obtained using the feature-table to ensure processing was successful. Taxonomy was assigned by training a naive Bayes classifier with the feature-classifier plugin [24], based on a 99% similarity to the V5-V6 region of the 16S rRNA gene in the SILVA 138 database to match the 784F/1061R primer pair used [27]. Mitochondria and chloroplast reads were filtered out. Analyses were performed using Rstudio version 2022.02.2 and the phyloseq package [28]. Metadata file, taxonomy table, phylogenetic tree and ASV table were imported into R to create a phyloseq object. Contaminant ASVs, arising from kit reagents and sample manipulation, were identified using the package decontam [29]. The function ‘isNotContaminant’ was used as it is more stringent and more adequate for low-biomass samples. One replicate for each OI2 and OI3 were removed because of heavy contamination. It is worth noting that bacteria belonging to the *Brachybacterium* genus accounted for more than 90% of the contamination (Table S1), which were contaminants of the Qiagen PCR kit previously observed [3]. For the re-analysis of previously acquired data, raw reads were downloaded from SRA (SRP150755 for X7, and SRP187380 for OI2 and OI3). Data was re-analyzed as described above. Plots were generated using GraphdPad Prism 9.

### Genome amplification of LCM samples and shotgun sequencing

To reach sufficient quantities for shotgun sequencing, DNA samples obtained through LCM were amplified as previously described [3], using a SeqPlex DNA Amplification Kit (Sigma-Aldrich, USA), which is specifically designed for low-quantity, fragmented DNA. The same DNA samples that were used for 16S rRNA gene metabarcoding were used and, for each sample, 1 µL (OI2 and OI3) or 3 µL (X7) of each of the three replicates were pooled before amplification. Only the X7 sample yielded sufficient DNA for sequencing. Samples were then sent to the Australian Genomic Research Facility (Melbourne, Australia) for sequencing. Library preparation was performed with an IDT xGen cfDNA & FFPE DNA Library Prep kit (Integrated DNA Technologies, USA) and samples were sequenced on a NovaSeq 6000 S4 2×150 bp Flowcell Illumina platform.

### Metagenomic data analysis

Quality control was performed using FastQC v0.11.9 [30]. Adapters, low-quality sequences (phred score < 30), as well as the first and last 10 bp were trimmed using Trim Galore v0.6.2 [31]. High-quality reads were mapped against a *P. acuta* draft genome [32] using Bowtie2 v2.4.2 with default parameters [33] to remove host-related reads from the data. Mapped reads were then removed using Samtools v1.11 [34]. Only paired-end host-removed reads were used for metagenome assembly. Metagenome assembly was carried out using MEGAHIT v1.2.9 [35] with a minimum contig length of 1000 bp and the following k-mers: 21, 33, 55, 77, 99. Contigs from kmer 99 were used for all downstream processing. Contig taxonomy was assessed using CAT/BAT v5.2.3 [36], and contigs assigned to Eukarya, Archea, or Virus were removed. Assembled contigs were binned using the binning (with metabat2, concoct and maxbin2 tools) and bin_refinement (with >70% completeness and <10% contamination as cut-off parameters) modules of MetaWRAP v1.3.2 [38]. Bin quality and taxonomy were assessed using CheckM v1.2.2 [39] and GTDB-Tk v2.3.0 [40], respectively. These bins were reassembled to further improve the contiguity and bin completeness and contamination stats using the reassemble_bins module implemented in MetaWRAP v1.3.2. The final taxonomy of individual contigs on a per bin level was assessed using CAT/BAT v5.2.3 [36], and any contig belonging to a different phylum than the taxonomy assigned by GTDB-Tk was manually removed. Bin coverage was obtained using CoverM v0.6.1 (https://github.com/wwood/CoverM) using the “genome” option. The full sequence of the 16S rRNA gene (1569 bp) was obtained using Barrnap v0.9 [41].

### Pure culturing of bacteria from *Pocillopora acuta*

Using clean bone cutters, fragments between 3 and 7 cm in length were sampled from the P3 colony, rinsed in FSW, and placed in sterile zip-lock bags containing FSW. Using a water flosser (Waterpik® Water Flosser) fitted with a clean nozzle, coral tissue was blasted off the skeleton. The resulting slurry was collected in a sterile zip-lock bag and transferred to 50 mL Falcon tubes. The tubes were centrifuged at 3750 × *g* for 10 min (Allegra X-12R, Beckman Coulter, USA) to obtain coral tissue pellets. The pellets were resuspended in 1 mL FSW and transferred to a single 1.5 mL tube per colony and centrifuged at 5000 × *g* for 10 min to remove any residual mucus from the sample. The tissue pellets were resuspended in 1 ml of FSW using a tissue lyser (Tissue-Lyser II, Qiagen, Australia) and the resuspension was homogenized using sterile glass homogenizers for 30 s. The resulting tissue homogenates were each serially diluted from 10^-1^ to 10^-7^ and plated in triplicates onto Marine Agar 2216 (MA, BD Difco). Following one week of incubation at 23 °C, individual bacterial colonies were selected and streaked onto fresh MA plates. Purified bacterial isolates were obtained after a further 2 rounds of sub-culturing onto fresh MA plates following a weekly incubation period at 25 °C and 26 °C, respectively.

### 16S rRNA gene sequencing of cultured bacteria

Individual, pure, freshly grown bacterial colonies were suspended in 20 μL sterile Milli-Q® water, incubated for 10 min at 95°C then used as templates in colony PCRs. PCR amplification of the bacterial 16S rRNA gene was with primers 27F and 1492R [42]. The PCR was performed with 20 µl Mango Mix™ (Bioline, UK), 0.25 µM of each primer and 2 µl of DNA template in a final volume of 40 µl with nuclease free water (Ambion, Thermo Fisher Scientific Inc., TX, USA). The thermal cycling protocol was as follows: 94°C for 5 min; 30 cycles of 94°C for 1 min, 50°C for 45 s and 72°C for 90 s; and a final extension of 10 min at 72°C. Amplicons were purified and sequenced on an ABI sequencing instrument by Macrogen (Seoul, South Korea). Trimmed high-quality read data from each isolate were used for presumptive identification by querying the 16S rRNA gene sequences via Blastn and compared to other 16S rRNA gene sequences obtained in this study.

### DNA extraction and whole-genome sequencing

The P3-11-1 isolate was selected for whole-genome sequencing. A single colony was picked with an inoculation loop and DNA extraction was performed on the QIAsymphony using the DSP Virus/Pathogen Mini Kit (Qiagen). Library preparation performed using Nextera XT (Illumina Inc.) according to manufacturer’s instructions. Whole-genome sequencing was performed on NextSeq 500/550 with a 150bp PE kit.

### Genome assembly, taxonomy, and annotation

Raw reads were trimmed and quality-filtered using Trimmomatic v0.36 [43] (HEADCROP:10 LEADING:5 TRAILING:5 SLIDINGWINDOW:4:28 MINLEN:30). The quality of raw reads before and after trimming was checked with FASTQC v0.11.9 [30]. Trimmed and quality-filtered reads were de novo assembled into contigs using SPAdes v3.15.5 [44] with 21, 33, 55, 77 and 99 k-mers, and the option “--careful” was applied to minimise the number of mismatches and short indels. All contigs with a length of <1000 bp were removed using BBMap v38.96 [45]. Subsequently, the levels of completeness and contamination of assembled genomes were assessed using CheckM v1.2.2 using the “lineage_wf” workflow [39]. Coverage was obtained using CoverM v0.6.1 (https://github.com/wwood/CoverM) using the “genome” option.

Taxonomic assignment of all assembled genomes was carried out using GTDB-Tk v2.3.0 [40] using the “classify_wf” workflow. GTDB-Tk assigned the taxonomy of the genomes based on 120 bacterial marker genes. Based on the GTDB-Tk alignment of the two genomes from this study and 28 additional genomes (Table S5), a phylogenetic tree was built in IQ-Tree v2.2.2.3 [46] using the best model Q.insect+F+R4, selected by ModelFinder wrapped in IQ-tree [47], and 1000 ultrafast bootstrap replicates [48]. The tree was visualized in iTOL v6 [49]. *Zookishella* was chosen as an outgroup. Average Nucleotide Identities (ANI) and Average Amino acid Identities (AAI) were calculated using a genome-based matrix calculator [50] and plotted in R using the pheatmap package [51]. Gene prediction was performed in Bakta v1.7.0 [52], KEGG-mapper Reconstruct [53], and InterProScan v5.55 with Pfam domain annotations [54]. Pfam annotations were used to look for eukaryotic-like proteins (ankyrin-repeat domains, WD40 domains, tetratricopeptide repeat) and the *dsyB* gene (PF00891). Secondary metabolites were predicted using antiSMASH v7.0.0 [55].

The 16S rRNA gene phylogenetic tree of the Endozoicomonadaceae family was constructed using 38 published, full-length 16S rRNA gene sequences from the Endozoicomonadaceae family. The full-length 16S rRNA gene (1572 bp) obtained from the P3-11-1 assembly was used for the phylogenetic tree construction. A MAFFT alignment was created using Geneious Prime v2019.1.3. The full alignment was stripped of columns containing 99% or more gaps. This alignment was used to generate a maximum likelihood phylogenetic tree with 1000 ultrafast bootstraps using IQ-Tree v2.2.2.3 [46] with the best model TIM3+F+I+G4, selected by ModelFinder wrapped in IQ-tree [47]. Sequences belonging to the *Zookishella* genus were chosen as an outgroup.

