## Supplemental material for "A novel, tissue-associated and vertically transmitted bacterial symbiont in the coral *Pocillopora acuta*"

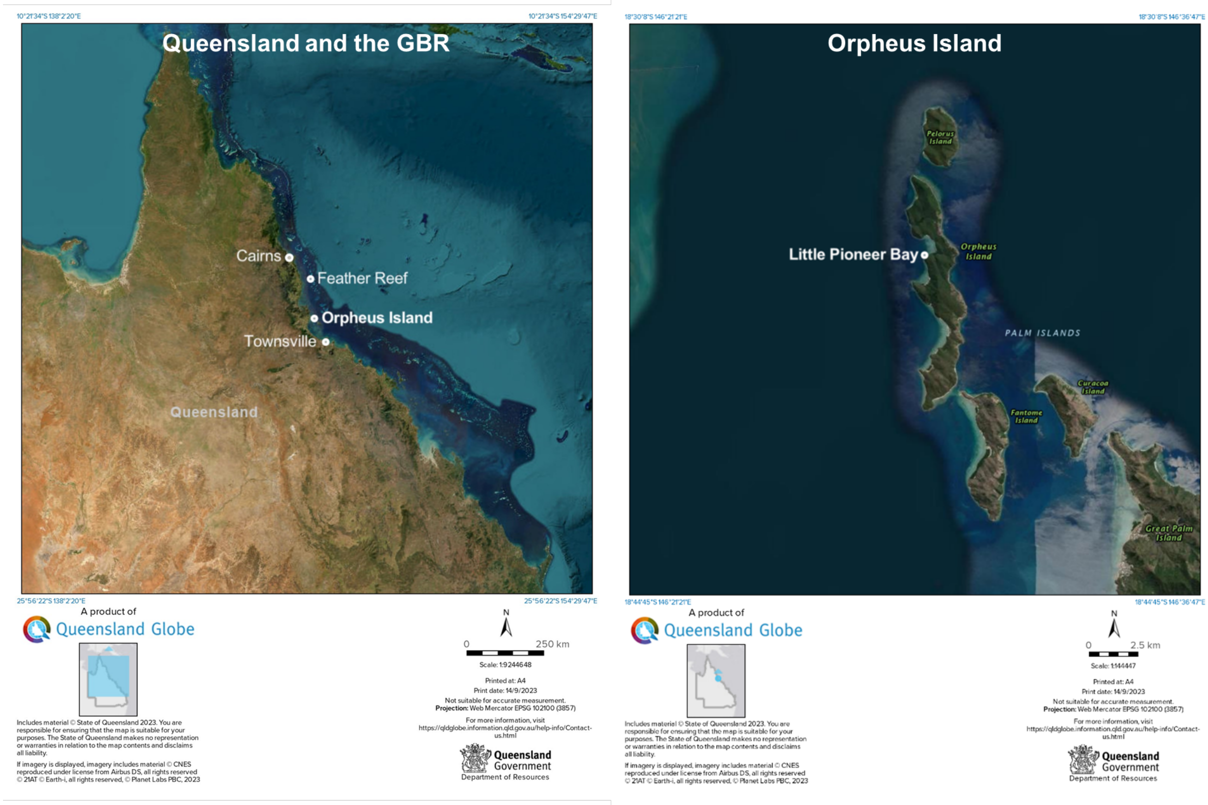


**Figure S1:** Map of study site at Little Pioneer Bay (Orpheus Island) in the central Great Barrier Reef. Feather Reef is also highlighted as it was the site of a previous study on CAMAs in *Pocillopora acuta* [1]. © The State of Queensland (Department of Resources) 2023

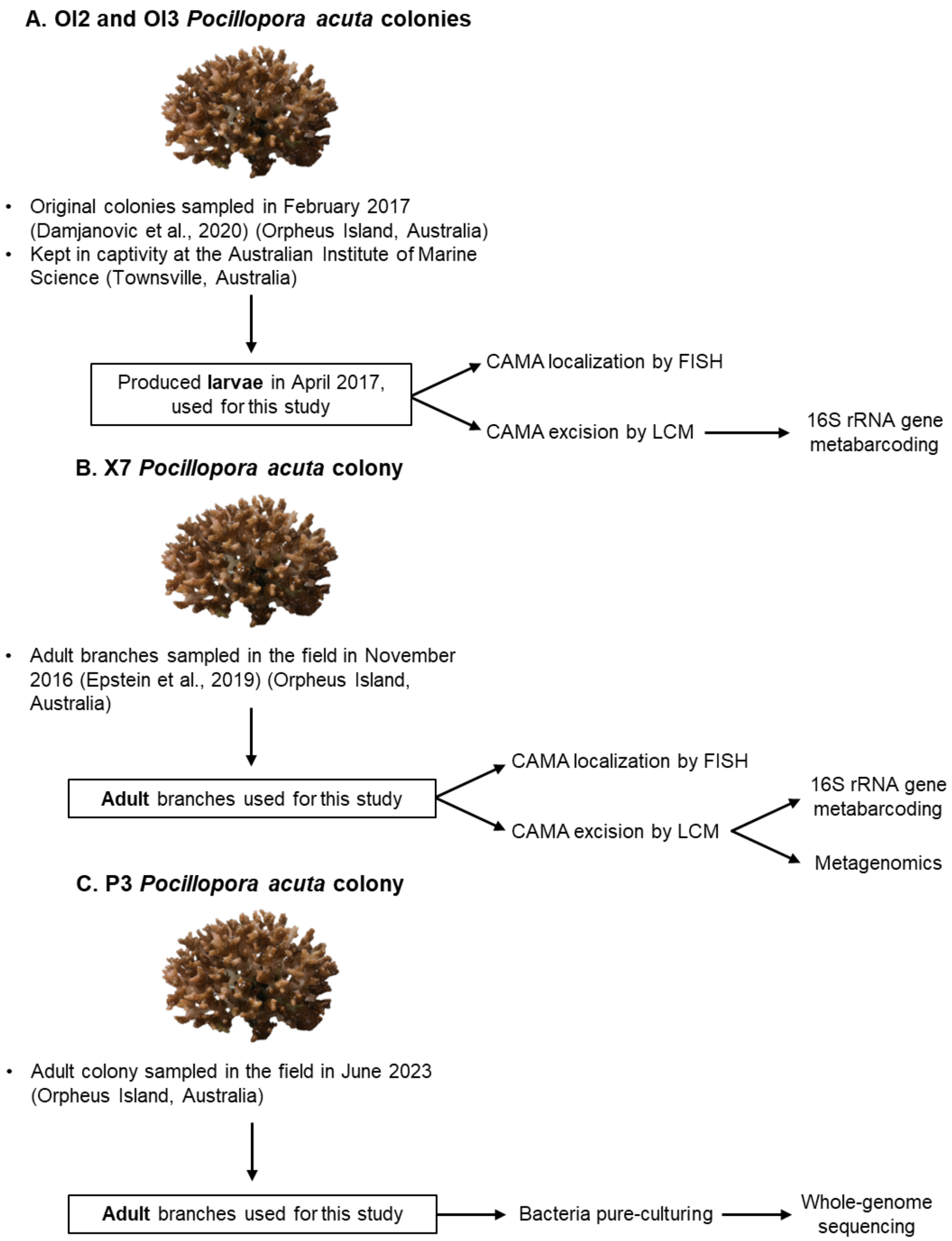


**Figure S2:** Sampling and experimental design used in this study. All *Pocillopora acuta* colonies were sampled from the same site in Little Pioneer Bay, Orpheus Island (Great Barrier Reef, Australia). A: Sampling of the OI2 and OI3 colonies (by Damjanovic et al. (2020) [1]). Following the establishment of the original colonies in captivity, larvae released by the adults were sampled. Larvae were used for microscopic imaging and 16S rRNA gene metabarcoding of CAMAs. In the original study, the adults and recruits were also sampled. B: Sampling of the X7 colony (by Epstein et al. (2019) [2]). Adult branches were sampled in the field and fixed. Branches were used for microscopic imaging, and 16S rRNA gene metabarcoding and metagenomics of CAMAs. C: Sampling of the P3 colony. Adult colonies were sampled in the field, and live fragments were used for the pure-culturing of bacteria and whole-genome sequencing of the isolated bacteria. FISH: fluorescence in situ hybridization; CAMA: cell-associated microbial aggregate; LCM: laser capture microdissection

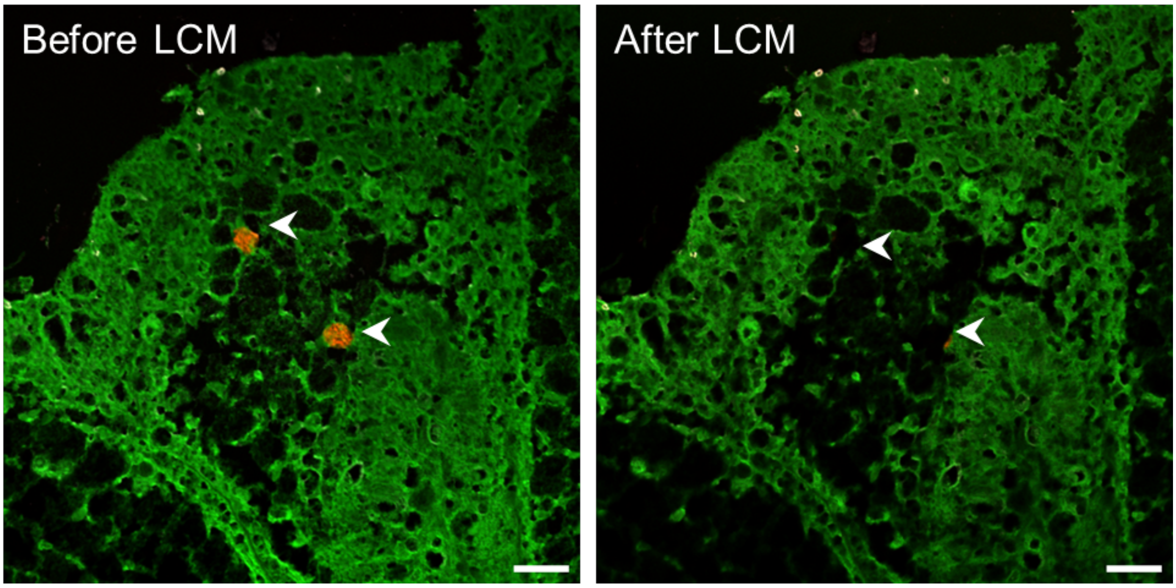


**Figure S3:** Laser capture microdissection (LCM) of CAMAs in *Pocillopora acuta*. Arrowheads point at CAMAs (left panel) and captured CAMAs (right panel) in the same section. Green: autofluorescence; red: EUB338-mix probe (all bacteria); white: non-EUB probe (negative control). Scale bars: 20 µm.

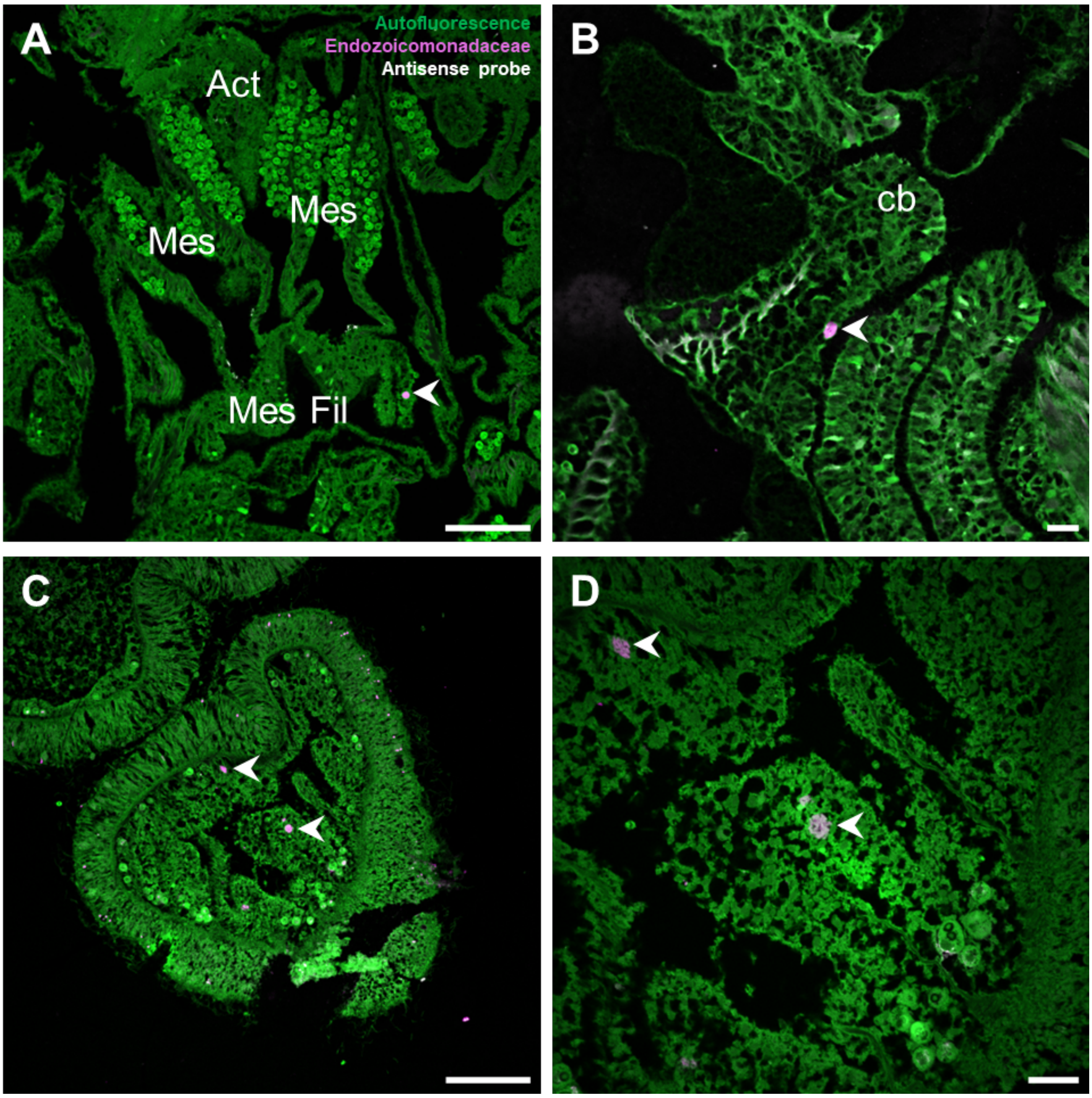


**Figure S4:** Endozoicomonadaceae in CAMAs of Orpheus Island *Pocillopora acuta*. CAMA location by FISH on sectioned adult polyps (A, B) and larvae (C, D). Arrowheads point at CAMAs. Green: autofluorescence; magenta: End663 probe (Endozoicomonadaceae); white: non-EUB probe (negative control). Act: actinopharynx; Mes Fil: mesenterial filaments; Mes: mesenteries; cb: cnidoglandular band. Scale bars: 100 µm for A, C; 20 µm for B, D.

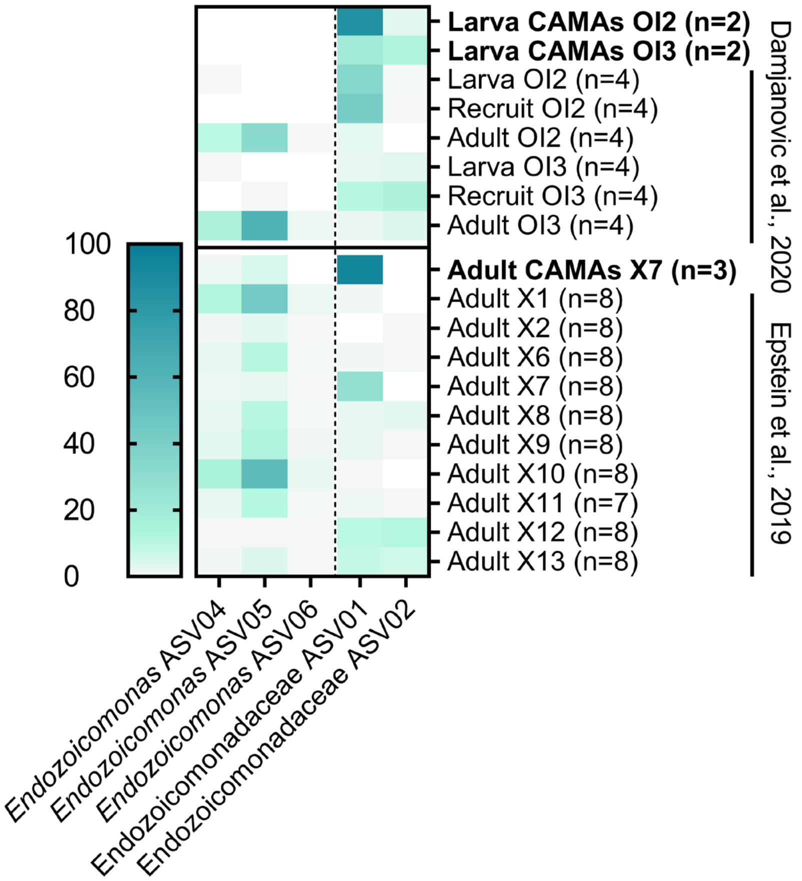


**Figure S5:** Relative abundance of the five most abundant Endozoicomonadaceae ASVs in CAMAs, whole larvae, recruits, and adults of Orpheus Island *Pocillopora acuta*. Rows in bold (CAMAs) are newly obtained data. Other rows represent reanalyzed data from Damjanovic et al., 2020 [2] and Epstein et al., 2019 [3]. Endozoicomonadaceae ASVs 01 and 02 were assigned as *Kistimonas* in our reanalysis, while they were initially assigned to *Endozoicomonas* in the original studies. Raw data are available in Table S2 (for CAMAs) and Table S3 (for the reanalysis of previous data).

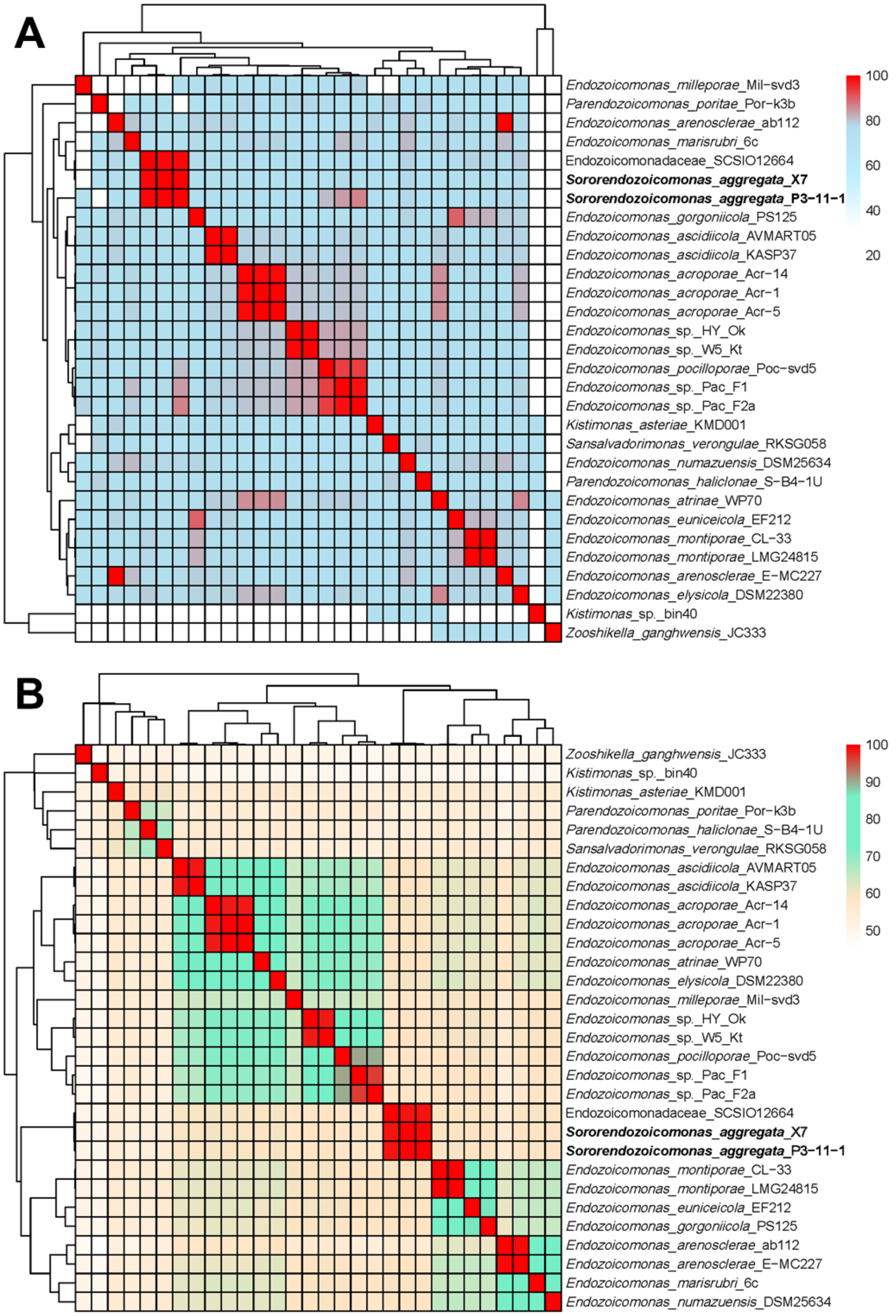


**Figure S6:** Average nucleotide identity (ANI) (A) and average amino acid identity (AAI) (B) of Pac_X7 and Pac_P3-11-1 with other Endozoicomonadaceae genomes. Additional data on the reference genomes are available in Table S5.

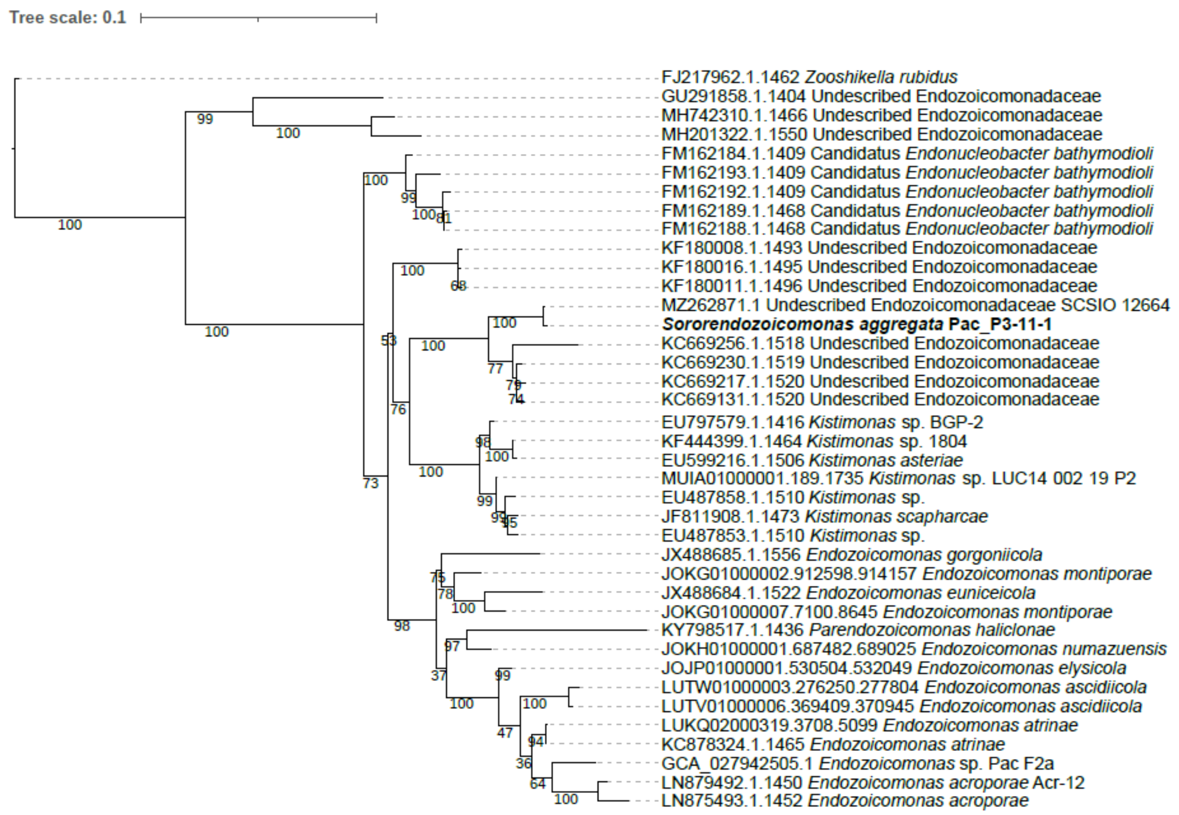


**Figure S7:** Maximum likelihood phylogenetic tree of 16S rRNA gene sequences showing the placement of Pac_P3-11-1 (in bold) in the Endozoicomonadaceae family based on 38 bacterial 16S rRNA sequences, in addition to the full 16S rRNA sequence obtained in this study. Bootstraps values based on 1000 replications are provided.

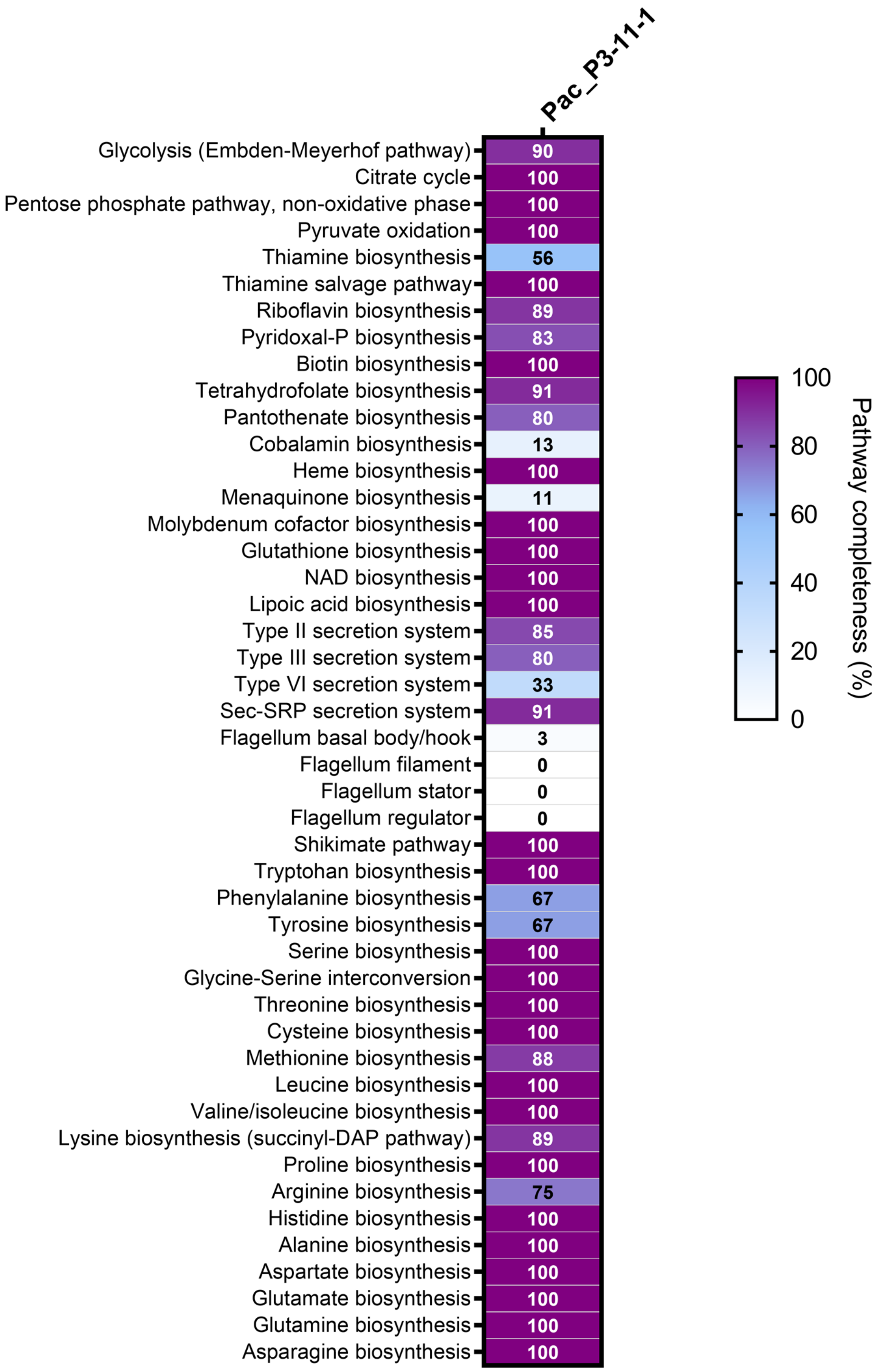


**Figure S8:** Estimated completeness of KEGG pathways of interest in the Pac_P3-11-1 genome recovered in this study.

**Table S1:** Sequencing statistics for the two 16S rRNA gene metabarcoding experiments analyzed in this study and shown in Figure 1F. The first column ("Larva CAMAs") refers to Table S2A, the second column ("Adult CAMAs") refers to Table S2B.

| **Experiment** | **Larva CAMAs (OI2 and OI3 colonies)** | **Adult CAMAs (X7 colony)** |
|---|---|---|
| **Total Samples (negative controls)** | 18 (12) | 11 (8) |
| **Raw reads** | 1323009 | 2426491 |
| **Reads after merging, denoising and chimera filtering** | 941134 | 1755498 |
| **Samples kept for analysis** | 4 | 3 |
| **ASVs after decontamination** | 18 | 12 |
| **Read per sample** | 13647 | 74591 |
| **Contamination (%)** | 82.9 | 29.4 |
| **Proportion of *Brachybacterium* sp. among contaminants (%)** | 93.2 | 89.8 |

**Table S2:** Relative abundance of bacterial ASVs in CAMAs isolated by LCM from larvae of the OI2 and OI3 colonies (A) and from adults of the X7 colony (B). Each column is a biological replicate. These data are summarized in Figure 1F.
(attached)

**Table S3:** Relative abundance of Endozoicomonadaceae ASVs in larvae, recruits, and adults of the OI2 and OI3 colonies (A), and adults of 10 colonies, including X7, from Orpheus Island based on a reanalysis of data reported in Damjanovic et al. (2020) [2] and Epstein et al. (2019) [3], respectively. Each column is the average of four (A) or seven to eight (B) biological replicates. These data are summarized in Figure S5.
(attached)

**Table S4:** Summary of the two assemblies recovered from bacterial pure cultures (Pac_P3-11-1) and from X7 CAMAs (Pac_X7)

| Genome | Pac_P3-11-1 | Pac_X7 (MAG) |
|---|---|---|
| **Size (bp)** | 4041637 | 3933347 |
| **Coverage (X)** | 61 | 37221 |
| **Completeness (%)** | 97.13 | 95.91 |
| **Contamination (%)** | 0.88 | 1.01 |
| **G+C content (%)** | 46.7 | 46.7 |
| **N50** | 37409 | 9295 |
| **L50** | 35 | 132 |
| **Number of contigs** | 201 | 605 |
| **Number of coding sequences (CDSs)** | 3545 | 3405 |
| **Number of ribosomal RNAs** | 4 | 3 |
| **Number of transfer RNAs** | 56 | 48 |

**Table S5:** List of Endozoicomonadaceae genomes used for phylogenetic analyses. (attached)

**Table S6:** Detailed Bakta annotations for the Pac_P3-11-1 genome recovered in this study. (attached)

**Table S7:** Number of eukaryotic-like protein sequences found in the Pac_P3-11-1 genome. Sequences were detected based on an InterProScan classification with Pfam domain annotations.

| **Genome** | **Pac_P3-11-1** |
|---|---|
| Ankyrin-repeat proteins | 12 |
| WD40 domain proteins | 14 |
| Tetratricopeptide repeat proteins | 9 |

**Table S8:** List of predicted secondary metabolites in the Pac_P3-11-1 genome recovered in this study.

| **Contig name** | **Type** | **Most similar known cluster** | **Biosynthetic gene cluster** | **Similarity (%)** | **Prokka annotation** |
|---|---|---|---|---|---|
| contig_6 | RiPP-like | - | - | - | AP-endonuc-2 domain-containing protein |
| contig_24 | betalactone | fengycin | BGC0001095 | 13% | Acetyl-CoA synthetase |
| contig_37 | NI-siderophore | fulvivirgamide A2/fulvivirgamide B2/fulvivirgamide B3/fulvivirgamide B4 | BGC0002620 | 66% | Siderophore synthetase component |

**Table S9:** List of oligonucleotides probes used for Fluorescence *in situ* Hybridization.

| Target group | Probe (fluorophore) | Sequence (5'-3') | % Formamide | NaCl concentration in washing buffer (M) | Ref |
|---|---|---|---|---|---|
| All bacteria | EUB338-mix (atto550) | GCWGCCWCCC GTAGGWGT | 25 | 0.149 | [4] |
| Negative control | nonEUB (atto647) | ACATCCTACG GGAGG | 25-35 | 0.07 - 0.149 | [5] |
| Endozoicomo ndaceae | End663 (atto550) | GGAAATTCCA CACTCCTC | 35 | 0.07 | [6] |
